# Inhibition of DNA-PKcs impairs the activation and cytotoxicity of CD4^+^ helper and CD8^+^ effector T cells

**DOI:** 10.1101/2022.06.23.497236

**Authors:** Ana C. Azevedo-Pouly, Lauren E. Appell, Lyle Burdine, Lora J. Rogers, Lauren C. Morehead, Melanie Barker, Zachary J. Waldrip, Brian Koss, Marie Schluterman Burdine

## Abstract

Modulation of T cell activity is an effective strategy for the treatment of autoimmune diseases, immune-related disorders and cancer. This highlights a critical need for continued investigation of proteins that regulate T cell function. The kinase <u>DNA</u>-dependent <u>p</u>rotein <u>k</u>inase <u>c</u>atalytic <u>s</u>ubunit (DNA-PKcs) is emerging as a potent regulator of the immune system spurring interest in its use as a therapeutic target for immune-related diseases. In murine models of autoimmune disease including asthma and rheumatoid arthritis, treatment with small molecule DNA-PKcs inhibitors, which are in clinical trials for cancer therapy, decreased disease severity. Additionally, DNA-PKcs inhibitors reduced T cell-mediated graft rejection and extended graft survival in a murine allogenic skin graft rejection model. These *in vivo* studies suggest the therapeutic use of DNA-PKcs inhibitors for autoimmune and T cell-mediated disorders. In this study, we sought to further characterize the effects of DNA-PKcs inhibitors on T cells to better understand their clinical potential. We determined that pharmacological inhibition of DNA-PKcs abrogated activation of murine and human CD4^+^ and CD8^+^ T cells as evident by reduced expression of the activation markers CD69 and CD25. Furthermore, inhibition of DNA-PKcs impeded metabolic pathways and proliferation of anti-CD3/CD28 activated CD4^+^ and CD8^+^ T cells as well as peptide-stimulated OTI-CD8^+^ T cells. This reduced the ability of OTI-CD8^+^ T cells to kill cancer cells and the expression of IFN*γ* and the cytotoxic genes eomes, perforin and granzyme B. These results suggest a novel role for DNA-PKcs in early T cell activation. Furthermore, our data support the therapeutic potential of DNA-PKcs inhibitors on diseases of immune dysregulation.

## Introduction

While T cells largely serving a beneficial role through elimination of diseased cells and tumor surveillance, dysregulated CD4^+^ and CD8^+^ T cells can be pathogenic. Autoimmune diseases including rheumatoid arthritis, multiple sclerosis, lupus, and asthma result from autoreactive T cells that release high levels of pro-inflammatory cytokines and chemokines and direct lysis of host cells.^1,2^ In addition, T cells drive allogenic organ rejection and acute graft versus host disease (aGVHD).^3,4^ Patients receiving donated organs must adhere to a life-long regimen of immunosuppressants to prevent rejection. Given the effects T cells have on health and disease, therapies designed to control their activity have an enormous impact clinically. Numerous small molecule or cell-based immunotherapeutics are FDA-approved for the treatment of immune-related diseases. For example, inhibitors of T cell activation including tacrolimus and the recently developed Janus kinase (JAK) inhibitors are successful drugs for the treatment of organ transplant rejection and rheumatologic disorders.^5,6^ Continued investigation of novel molecules that regulate CD4^+^ and CD8^+^ T cell function is imperative for the generation of additional therapeutic options for life threatening autoimmune and immune-related diseases.

DNA-PKcs is a 460 kDa phosphatidylinositol 3-kinase-related (PIKK) serine/threonine kinase. In its canonical role, DNA-PKcs functions as a DNA damage repair kinase recruited to DNA double-strand break foci, whereupon activation via autophosphorylation of serine 2056, it induces a phosphorylation cascade that recruits repair proteins to initiate non-homologous end joining (NHEJ).^7,8^ DNA-PKcs inhibitors, highly-specific, synthetic small molecule ATP-competitors including NU7441, sensitize tumor cells to radiation and chemotherapy by reducing DNA damage repair resulting in decreased tumor growth in *in vivo* mouse models.^9–11^ Therefore, DNA-PKcs inhibitors are currently being evaluated as antineoplastic adjuvants in combination with radiation, chemotherapy and immunotherapy in Phase I/II clinical trials.^12,13^ Interestingly, DNA-PKcs is emerging as a critical regulator of the immune system. While it is known that DNA-PKcs is required for lymphocyte maturation and immune system diversity in developing mammals due to its role in V(D)J recombination and DNA repair, recent studies identified additional functions for DNA-PKcs in both the mature innate and adaptive immune responses independent of its role in DNA repair.^14–16^ For instance, DNA-PKcs is required for Toll-like receptor 9 (TRL9) signaling and interferon (IFN) *αβ* production in dendritic cells.^17^ DNA-PKcs residing in the cytoplasm acts as a viral DNA sensor inducing activation of the lRF3-dependent innate immune response. Inhibition of DNA-PKcs attenuated IFN-I responses and increased viral load.^18^ Mirsha et al. reported that in an murine asthma model, dust mite antigens activate DNA-PKcs in dendritic cells which is required for efficient antigen presentation to induce a Th2-mediated inflammatory response.^19^ Additionally, Ghonim et al. determined in a similar asthma model that DNA-PKcs regulated differentiation of CD4^+^ T cells into Th1 and Th2 subtypes by altering expression of the transcription factors Gata3 and Tbet.^20^ In both studies, treatment with DNA-PKcs inhibitors reduced the severity of asthma-related symptoms. In our previous studies, we determined DNA-PKcs is required for the activity of the transcription factors NFAT, NF*κ*B and EGR1 and production of cytokines critical to T cell function, including IL2, IL6, IFN*γ*, and TNFα.^21–23^ Moreover, we showed that treatment with the DNA-PKcs inhibitor NU7441 extended the survival of transplanted allogenic skin grafts in a murine mouse model by reducing T cell-graft infiltration and inflammatory cytokine production.^22^ These studies highlight the immunosuppressive effects of DNA-PKcs inhibitors and suggest the potential repurposing of DNA-PKcs inhibitors for immunosuppression therapy for T cell-mediated autoimmune and immune-related diseases and allogenic graft rejection. However, the effects of DNA-PKcs inhibition on T cells have not been thoroughly investigated but is critical for understanding potential clinical benefits. Therefore, in this study, we sought to further understand the effects of DNA-PKcs inhibition on T cells. We determined that treatment with NU7441 significantly disrupted the activation, metabolic pathways and cytotoxic function of stimulated CD4^+^ and CD8^+^ T cells. These data suggest a novel role for DNA-PKcs in early T cell activation and the immunosuppressant properties of DNA-PKcs inhibitors. Importantly, they highlight the need for clinical precautions when using DNA-PKcs inhibitors for cancer therapy and further support the potential use of DNA-PKcs inhibitors as therapy for immune-related disorders including aGVHD and organ transplant rejection.

## Results

### DNA-PKcs expression is elevated following TCR-mediated T cell activation

To confirm a function for DNA-PKcs in T cells, we first validated expression of DNA-PKcs in primary splenic CD4^+^ and CD8^+^ T cells by intracellular flow cytometry. Twenty-four hours following anti-CD3/CD28 activation, total DNA-PKcs expression significantly increased compared to unactivated levels (**Figure 1**). This observation was also observed by western blot where treatment with the DNA-PKcs inhibitor reduced DNA-PKcs expression following activation in CD8^+^ T cells. This result corresponds to our previous data indicating an increase in DNA-PKcs levels in human Jurkat T cells upon T cell activation and highlights a function for DNA-PKcs in stimulated primary T cells.^21^

**Figure 1.**
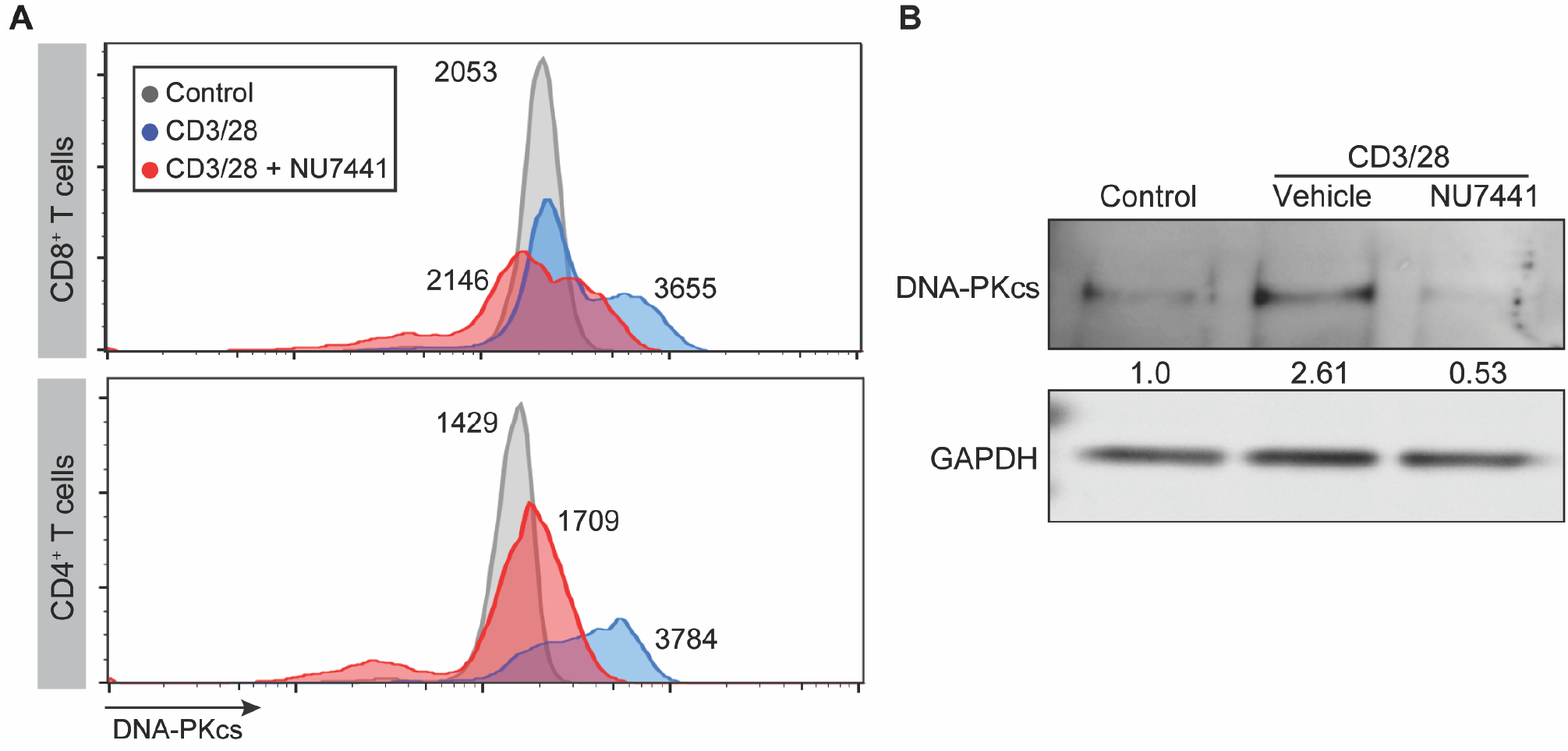
Elevated DNA-PKcs expression in activated CD4^+^ and CD8^+^ T cells. CD8+ or CD4+ T cells were activated (*α*CD3/CD28) in vehicle (DMSO) or 2 *µ*M NU7441 for 24hrs. (**A**) Flow cytometric analysis of intracellular DNA-PKcs was performed. Representative histograms of DNA-PKcs AF647 fluorescence in the three treatment groups is shown. Fluorescence intensity goes up following T cell activation and is reduced with the addition of the inhibitor. (**B**) Western blot of total DNA-PKcs expression in cells with no activation, activation, or activation plus inhibitor. N=3 (pooled).

### Inhibition of DNA-PKcs impairs activation of CD4^+^ and CD8^+^ T cells

Our previous studies determined that treatment with the DNA-PKcs inhibitor NU7441 reduced inflammatory cytokine production by T cells and reduced proliferation induced by alloantigen recognition.^21,22^ To gain better insight into these results, we assessed the effect of NU7441 on T cell activation. Splenic CD4^+^ T cells isolated from wildtype mice were stimulated with anti-CD3/CD28 antibodies and monitored for activation via expression of two prominent cell surface activation markers, CD69 and CD25. Activation elevated membrane expression of both CD69 and CD25 in CD4^+^ T cells. However, inhibition of DNA-PKcs via NU7441 treatment prevented the increase in double positive CD69 and CD25 cells by 73% at 24hrs and by 95% at 48hrs compared to untreated activated cells (**Figure 2A**). Similar results were also seen in isolated primary CD8^+^ T cells where NU7441 treatment reduced CD69 and CD25 expression by 97% at 24hrs and 92% at 48hrs (**Figure 2B**). These results suggest DNA-PKcs inhibition abrogates T cell receptor (TCR)-induced activation of CD4^+^ and CD8^+^ T cells.

**Figure 2.**
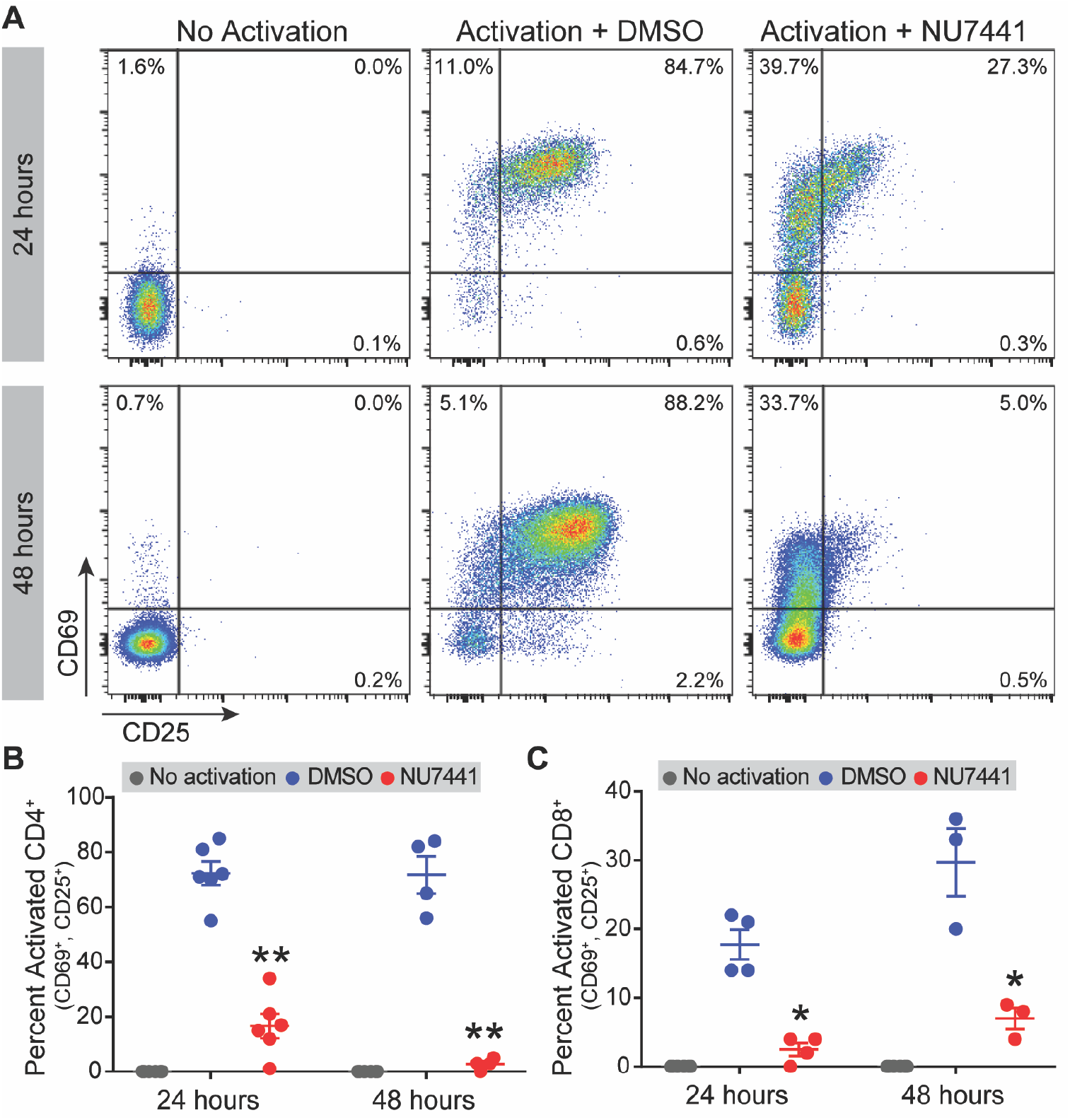
Inhibition of DNA-PKcs impairs CD4^+^ and CD8^+^ T cell activation. CD4^+^ (**A**,**B**) or CD8^+^ (**C**) T cells were activated (*α*CD3/CD28) in the presence of vehicle (DMSO) or 2 *µ*M NU7441 for 24 and 48 hrs. Flow cytometric analysis of CD25 and CD69 activation markers (viability 7-AAD) expression was then performed. (**A**) Representative staining for each CD4+ experimental group. Error bars S.D. of n=3 or more. *p ≤ 0.05, **p.001.

### Metabolism and proliferation in activated T cells are altered by DNA-PKcs inhibition

Immediately following TCR-induced activation through engagement of cognate peptide-MHC complexes, CD4^+^ and CD8^+^ T cells undergo metabolic changes to accommodate for increased cellular proliferation and function.^24^ These changes signal T cell activation and include an increase in glucose uptake and aerobic glycolysis. To further confirm a defect in early T cell activation with DNA-PKcs inhibition, we monitored metabolic changes in CD4^+^ and CD8^+^ T cells activated with anti-CD3/CD28 in the presence of NU7441 by measuring the extracellular acidification rate (ECAR), an indicator of aerobic glycolysis. As expected, T cell activation profoundly induced ECAR indicating increased glycolytic metabolism (**Figure 3A**). However, inhibition of DNA-PKcs with NU7441 treatment prevented aerobic glycolysis engagement and reduced maximal ECAR levels in both CD4^+^ and CD8^+^ T cells. To analyze metabolic effects in CD8^+^ T cells activated with a true antigen, we employed the OVA-specific TCR transgenic OTI mouse model. OTI mice express a TCR designed to recognize the OVA class I antigen SIINFKEL.^25,26^ OTI-CD8^+^ T cells were activated with the SIINFKEL peptide and ECAR measured (**Figure 3B**). Inhibition of DNA-PKcs activity prevented aerobic glycolysis engagement. Additionally, NU7441 reduced proliferation of stimulated OTI-CD8^+^ T cells (**Figure 3C**). These data are consistent with an abrogation in T cell activation observed with DNA-PKcs inhibitor treatment.

**Figure 3.**
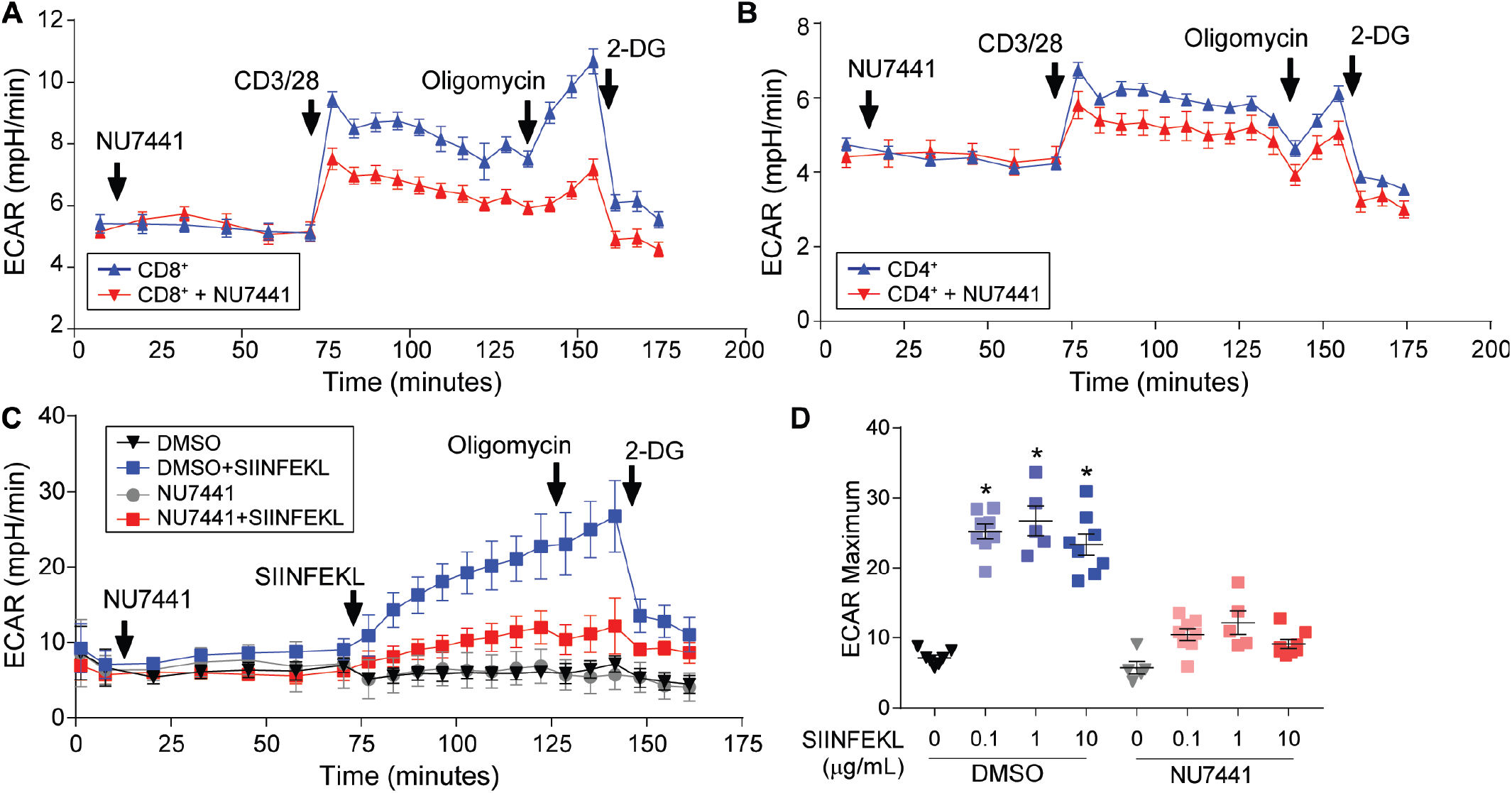
Inhibition of DNA-PKcs prevents engagement of aerobic glycolysis following activation regardless of antigen concentration. Representative trace of **(A)** CD8+ **(B)** CD4+ T cells stimulated with *α*CD3/CD28 (1 *µ*g/mL) while being monitored by a Seahorse XF96 metabolic flux analyzer. T cells treated with NU7441 (2*µ*M) one hour prior to stimulation had decreased glycolytic rate (ECAR). Oligomycin (ATP synthase inhibitor) and 2-DG (glycolysis inhibitor) were used as controls. **(C)** Representative trace of OTI-CD8+ T cells stimulated with the SIINFEKL peptide (1*µ*g/mL) while being monitored by a Seahorse XF96 metabolic flux analyzer as described above. **(D)** Maximal ECAR of T cells activated for 48hrs with different concentrations of SIINFEKL peptide. Error bars = S.D., **p≤0.01.

DNA-PKcs phosphorylates the serine/threonine kinase AKT at serine 473 resulting in kinase activation.^23,27^ AKT is phosphorylated in T cells following stimulation. This is required for downstream signaling which promotes the shift to aerobic glycolysis to support increased proliferation and gene transcription in activated T cells.^24,28^ We show that treatment with DNA-PKcs inhibitors significantly reduced AKT serine 473 phosphorylation highlighting a potential mechanism by which inhibitors disrupt T cell activation and metabolism (**Figure 4**).

**Figure 4.**
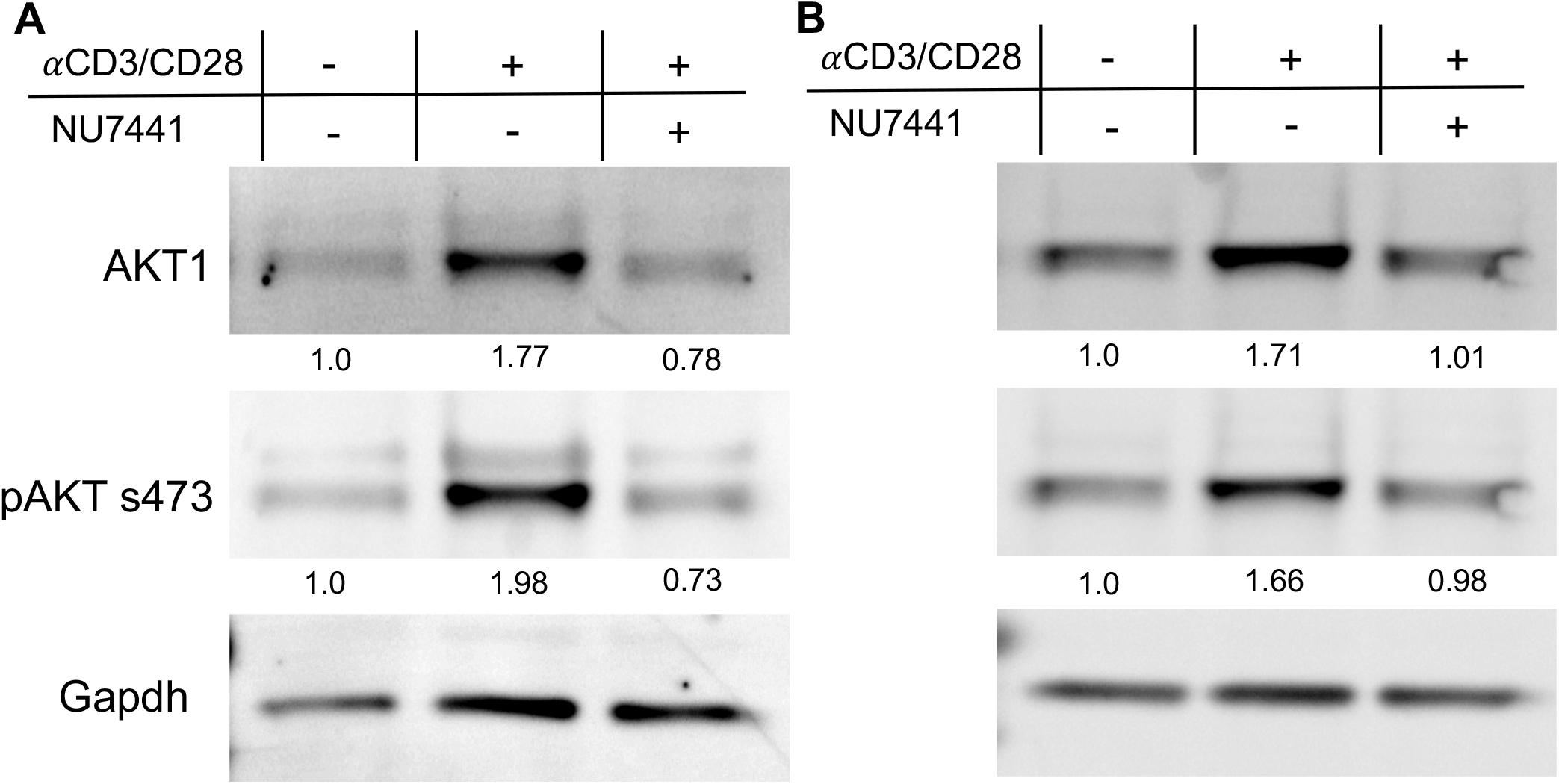
AKT protein expression is modulated by NU7441. Isolated mouse **A**) CD4^+^ or **B**) CD8^+^ T cells were activated (*α*CD3/CD28) in the presence of vehicle (DMSO) or 2*µ*M NU7441 for 24hrs. Western blot and normalized expression of AKT1 and pAKT to Gapdh for the three treatment groups is shown. N=3 (pooled).

### OTI-CD8^+^ T cells have impaired cytotoxicity following DNA-PKcs inhibition

We next sought to determine if the defects in T cell activation and metabolism mediated by DNA-PKcs inhibition translated into functional deficits. To do so, we analyzed the response of cytotoxic OTI-CD8+ T cells to SIINFEKL-expressing tumor cells. OTI-CD8^+^ T cells were incubated with SIINFKEL-MC38 colon cancer cells for 24 hours and viability of tumor cells analyzed. This study revealed that while untreated OTI-CD8^+^ T cells effectively killed the tumor cells at various effector-target ratios, NU7441 treated T cells had a significant decrease in cytotoxic function (**Figure 5A**). Ensuing activation, cytotoxic CD8^+^ T cells generate proteases and cytolytic proteins that mediated target cell killing. Such molecules include the transcription factor eomes, granzyme B and perforin as well as the inflammatory cytokine Interferon *γ* (IFN *γ*). DNA-PKcs inhibition significantly reduced expression of these proteins by activated OTI-CD8^+^ T cells consistent with a decrease in cytotoxicity and activation (**Figure 5B**).

**Figure 5.**
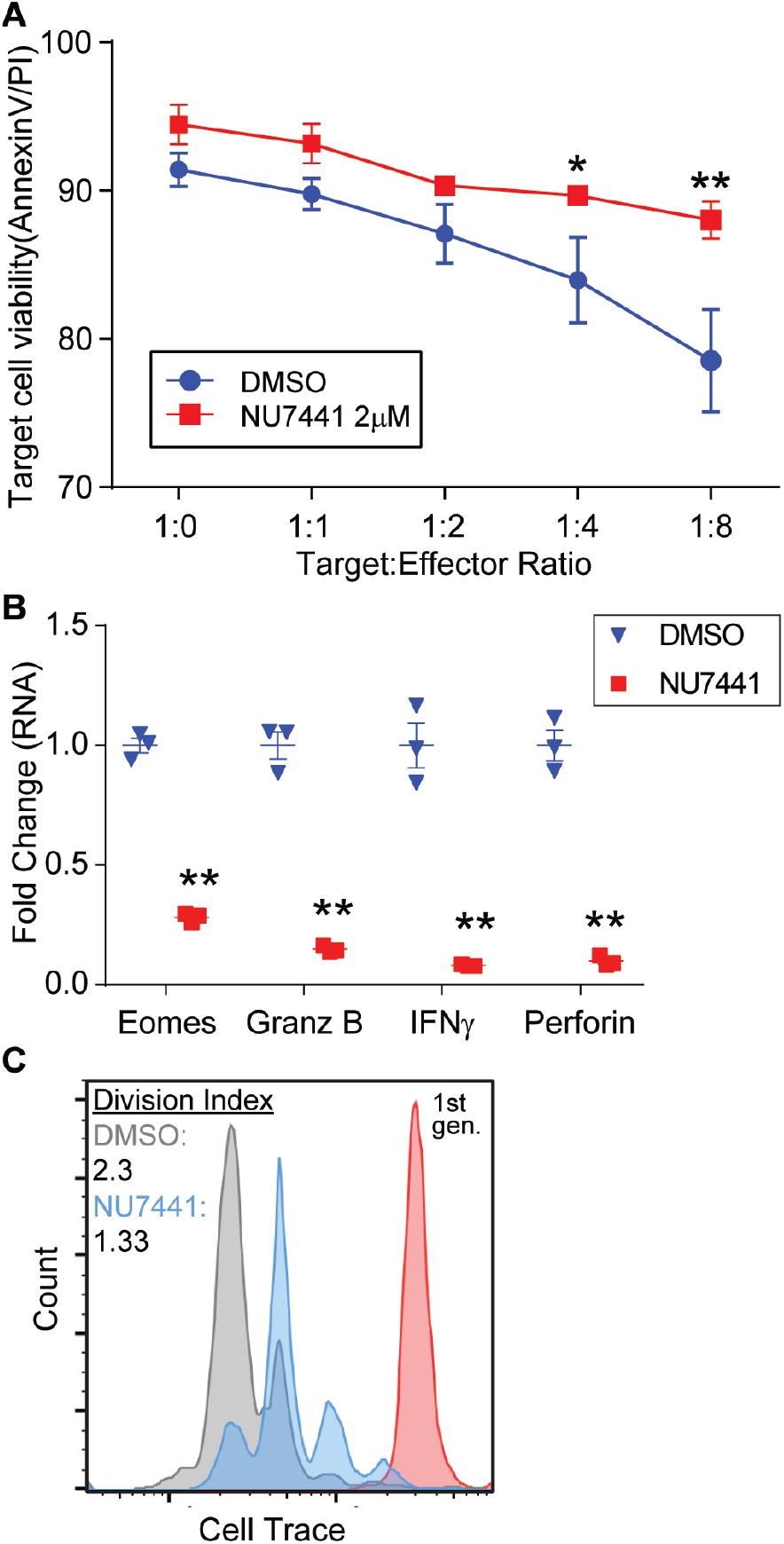
Loss of DNA-PKcs activity inhibits cytotoxicity of CD8^+^ T cells. OT-I CD8^+^ T cells were activated (*α*CD3/CD28) in the presence of vehicle (DMSO) or 2□M NU7441 for 48 hours. **(A)** Activated T cells were then co-cultured at increasing ratios with MC38-SIINFEKL cell line for 24 hours. T cells were stained with Cell Trace and viability of target cells was determined using Annexin V/PI staining. Error bars = S.D. of 3 independent experiments. **(B)** Cytotoxic gene expression at 48 hrs was determined using qPCR. Error bars represent the S.D. of n=3 independent experiments. *p ≤ 0.05, **p.001 **(C)** Representative cell division data of NU7441 (2□M) treated T cells. Prior to activation OTI CD8^+^ T cells were labeled with Cell Trace Violet dye and then activated with 1□g/mL SIINFEKL peptide for 48hrs. FACS analysis and Flow Jo software was used to calculate the division index.

### Inhibition of DNA-PKcs reduces activation of primary human T cells

Lastly, to provide relevance to human disease, we evaluated the effects of DNA-PKcs inhibition on human CD4^+^ and CD8^+^ T cells. T cells isolated from donated human PBMCs were activated with *α*CD3/CD28 and monitored for changes in activation, metabolism, and proliferation with or without NU7441 treatment. As seen in the above experiments using murine T cells, DNA-PKcs inhibitors decreased expression of the early and late activation markers CD69 and CD25 in human T cells 24 and 48 hours after *α*CD3/CD28 stimulation (**Figure 6A**). Furthermore, NU7441 disrupted metabolic pathways in human T cells preventing the induction of aerobic glycolysis following activation. We observed a reduction in maximal ECAR levels and aerobic glycolysis engagement in NU7441 treated-T cells compared to activated T cells without NU7441 treatment (**Figure 6B**). Taken together, these data confirm that DNA-PKcs inhibitors have a significant negative impact on T cell function and support previously published *in vivo* findings that DNA-PKcs inhibitors have immunosuppressant properties.

**Figure 6.**
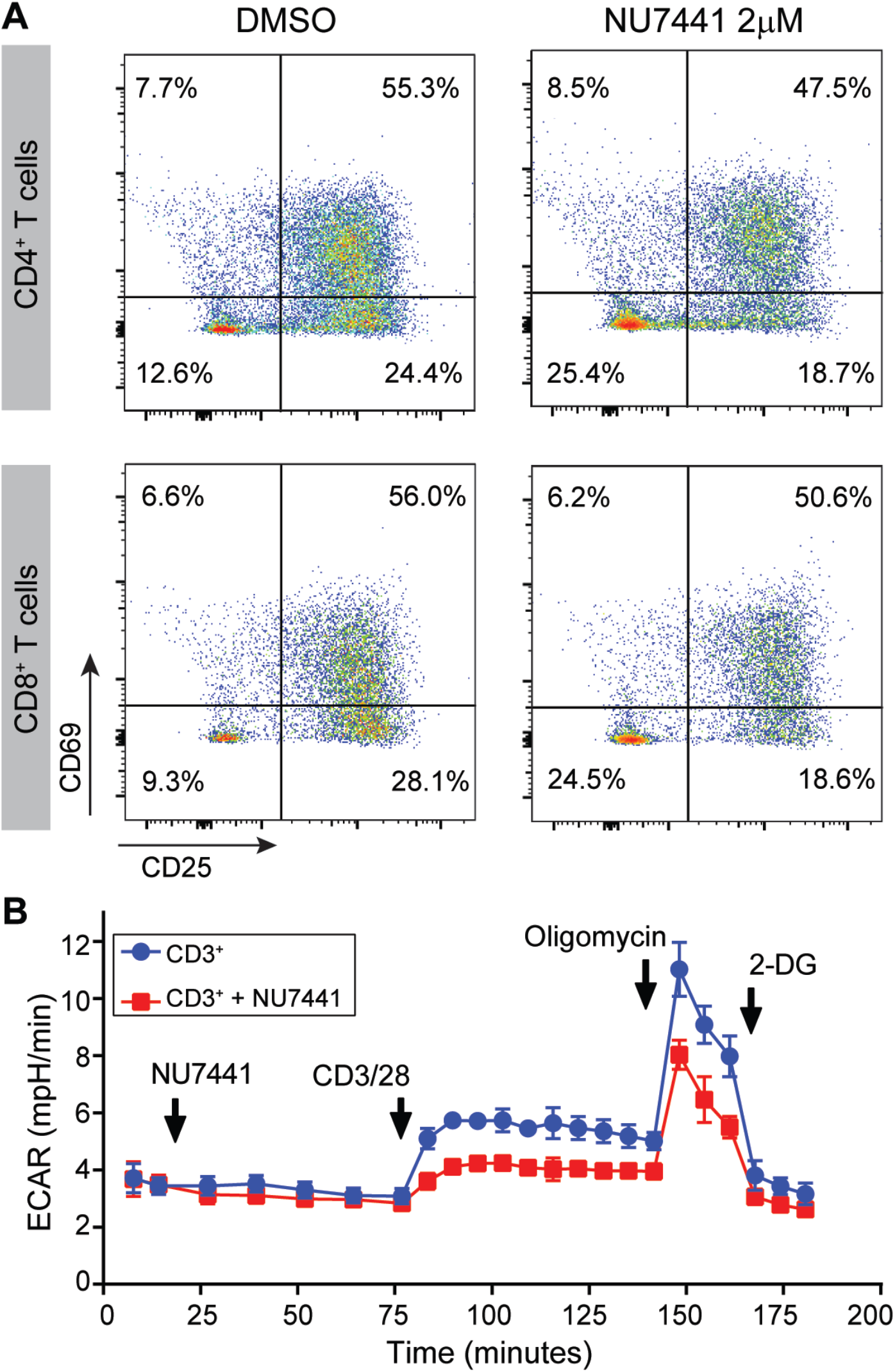
Inhibition of DNA-PKcs reduces activation of human T cells. **(A)** Representative expression of activation markers (CD25 and CD69) of human CD4^+^ and CD8^+^ T cells 48 hours post *α*CD3/CD28 mediated activation. **(B)** Representative trace of CD3^+^ T cells stimulated *α*CD3/CD28 while being monitored by a Seahorse XF96 metabolic flux analyzer. T cells treated with NU7441 (2*µ*M) one hour prior to stimulation had decreased glycolytic rate (ECAR). Oligomycin (ATP synthase inhibitor) and 2-DG (glycolysis inhibitor) were used as controls.

## Discussion

Substantial breakthroughs in the mechanistic understanding of T cell response to antigen and disease pathogenesis have made T cell modulation a promising area of therapeutic development. The success of cytokine-targeted therapy, particularly IL2 (Tacrolimus) and IL6 (Tocilizumab), and kinase inhibitors (JAK1 and 2) in the treatment of autoimmune diseases further deepens interest in this strategy.^5,6,29,30^ The re-evaluation of currently available drugs for alternative uses is a promising and cost-effective method for rapidly developing new treatments options. Small molecule inhibitors for the kinase DNA-PKcs were developed for the treatment of cancer given that the loss of DNA-PKcs activity results in the accrual of irradiation-induced DNA-damage in tumor cells and decreased tumor growth.^31^ However, recent studies highlighting effects of DNA-PKcs inhibitors on innate and adaptive immune cells suggest their potential use as therapy for autoimmune and immune-related diseases. While previous findings support an immunosuppressive function of DNA-PKcs inhibitors, their effects on T cells have not been thoroughly examined. Therefore, we set out in this study to characterize the effects of the DNA-PKcs inhibitor NU7441 on T cells in order to understand their potential use as immunosuppression therapy.

Here, we analyzed the effects of the DNA-PKcs inhibitor NU7441 on TCR-mediated activation, metabolism, proliferation and cytotoxicity using *ex vivo* cultured isolated murine and human CD4^+^ and CD8^+^ T cells. Our findings demonstrate that inhibition of DNA-PKcs has a significant effect on T cell activation evident by a decrease in the expression of early and late activation markers CD69 and CD25 following stimulation with anti-CD3/CD28 antibodies. This data highlights for the first time that DNA-PKcs may be involved in transduction of early TCR signaling induced by MHC-antigen/TCR engagement. In support of activation deficiencies, T cells treated with NU7441 also exhibited a defect in the switch to aerobic glycolysis normally induced by activation. In addition, we show that NU7441 treatment decreased proliferation of antigen stimulated OT-I CD8^+^ T cells and diminished the ability of T cells to lyse antigen-specific tumor cells in co-culture assays. These data indicate that DNA-PKcs inhibition has a significant effect on the activation and function of T cells and aligns with *in vivo* studies by Ghomin et al. and Mishra et al. where loss of DNA-PKcs activity by chemical inhibition resulted in decreased T cell and dendritic cell response to asthma-antigen exposure and highlighted a role for DNA-PKcs in Th2 immunity.^19,20^ In addition, these data are further supported by our previous publication where DNA-PKcs inhibition reduced T cell cytokine production and T cell migration into allogeneic skin grafts.^22^ However, in slight contrast to our studies, Tsai et al. reported that treatment with DNA-PKcs inhibitors increased expression of CD25 and the activation marker CD44 (CD69 was not analyzed) in CD4^+^ and CD8^+^ T cells.^32^ While this difference could be a result of an alternate activation method (anti-CD3 vs. anti-CD3/CD28 or OVA peptide), additional data from the report indicates that treatment with DNA-PKcs inhibitors alone reduced T cell proliferation and the infiltration of T cells into tumors. This supports the idea that DNA-PKcs inhibition suppresses T cell activation/function. Taken together, the above data confirm that DNA-PKcs inhibitors have immunosuppressive effects on T cells. This side effect should be considered when using DNA-PKcs inhibitors for cancer therapy where T cell activity is important for tumor control and may account for disappointing outcomes in current clinical trials. Importantly, it highlights the potential repurposing of DNA-PKcs inhibitors for immunosuppression therapy for T cell-mediated autoimmune diseases and allogenic graft rejection.

In the present study, we report one potential mechanism by which DNA-PKcs inhibitors curtail T cell activation and function is through reduction of AKT phosphorylation, a known target of DNA-PKcs. Activated AKT phosphorylates multiple downstream signaling molecules important for T cell activities including proliferation, glucose metabolism and differentiation.^33^ Continued investigation into additional mechanisms by which DNA-PKcs specifically is regulating T cell activation are currently underway in our laboratory. While DNA-PKcs expressed within the nucleus is known to bind to DNA to regulate DNA repair and gene transcription, it is interesting to note that DNA-PKcs is also expressed in the cytoplasm of numerous cell types including T cells and is involved in multiple pathways including DNA sensing, migration and endosomal signaling.^34–36^ Little is known about the role of cytoplasmic DNA-PKcs in T cells but further investigation could shed light on novel mechanisms that regulate T cell activation. For instance, DNA-PKcs is found localized to lipid rafts in the plasma membranes of irradiated cells and required for irradiation-induced intracellular signaling.^37^ In T cells, proteins that form the immune synapse following TCR-MHC/antigen engagement are localized to plasma membrane lipid rafts.^38^ It is an intriguing possibility that upon T cell stimulation, DNA-PKcs is localized to the immune synapses within lipid rafts where it impacts downstream signaling pathways that drive activation, metabolism and function. Additional investigations into the role of DNA-PKcs in immune cells will provide new insights into T cell activation that will uncover novel targets for therapeutic strategies.

As studies focused on DNA-PKcs in the immune system continue, it is clear that it is a key regulator of multiple aspects of immune cell function. It is imperative that the effects of DNA-PKcs inhibitors on immune cells are thoroughly described to assess their clinical use.

## Methods and Materials

### Reagents and antibodies

Flow cytometric antibodies – anti-CD3 eFlour 506, anti-CD4 eFlour 450, anti-CD8α eFlour 450, goat anti-rabbitt IgG (Invitrogen, #69-0032-80, 48-004-82, 48-0081-82, A32733); anti-CD25 FITC and anti-CD69 APC/Cy7 antibodies (BioLegend, #102006, 104526); anti-DNA-PKcs (Cell Signaling, #38168). Western blot antibodies - GAPDH antibody (ThermoFisher cat# MA5-15738); pAKT 473 (Cell Signaling, #4060), AKT (Cell Signaling, #2938). Inhibitors - NU7441 (Selleckchem, #S2638). T cell activation antibodies– purified anti-CD3ε and purified anti-CD28 (BioLegend, #100340, 102116). For all experiments, supplemented RPMI (RPMI-1640 supplemented with 10% fetal bovine serum, penicillin-streptomycin, Sodium Pyruvate, MEM NEAA, 30U/ml recombinant IL2 (ThermoFisher) and 55 *µ*M β-mercaptoethanol) was used.

### Mice

C57Bl/6 and OT-I mice were housed and bread in specific pathogen-free conditions at Arkansas Children’s Research Institute and the University of Arkansas for Medical Sciences. All animal studies were approved by the Institutional Animal Care and Use Committee of the University of Arkansas for Medical Sciences/Arkansas Children’s Research Institute.

### CD3ε^+^, CD8α^+^, and CD4^+^ T cell isolation

Single-cell suspensions enriched for naive CD3ε^+^, CD8α^+^ or CD4^+^ T cells from mouse spleens were isolated. Briefly, spleens were minced and passed through at 70 μm nylon cell strainer into PBS. After washing in PBS, CD8+ T cells were isolated from mouse splenocytes using CD3ε^+,^ Naïve CD4^+^ T or CD8α^+^ MACS T cell isolation kits (Miltenyi Biotech, #130-094-973, 130-104-453, 130-104-075) according to the manufacturer’s protocol. For in vitro co-culture experiments OTI+ T cells were purified using CD8α^+^ enrichment as detailed above.

### Primary cell culture

#### Murine T cell culture

CD3ε^+^, CD4^+^ and CD8α^+^ T cells were cultured as follows: plates were coated with 5 µg/ml anti-CD3ε (or with PBS alone) for 12-18 hrs at 4 °C. Cells were then plated at 1-2 × 10^6^ cells/ml in supplemented RPMI and either left unstimulated, stimulated with the plate-bound anti-CD3ε and 5 µg/ml of soluble anti-CD28, or stimulated and treated with 2-5µM of NU74411 for 24-48 hours.

#### Human T cell culture

CD3ε^+^ T cells were isolated using negative selection Human T cell Isolation Kit (StemCell Technology, #17951). Cells were plated at 1 × 10^6^ cells/ml in T cell growth media (StemCell Technology, # 10981) and activated using ImmunoCult™ Human CD3/CD28/CD2 T Cell Activator (StemCell Technology, #10970) with or without 2µM of NU74411 for 48 hours.

### Flow cytometry

Isolated CD3^+^, CD4^+^ and CD8α^+^ T cells were stained after activation and flow cytometry was performed. Cells were stained for cell surface markers and resuspended in eBioscinece flow cytometry staining buffer (Invitrogen, #00-4222-26). Viability was assessed by staining with 7-AAD Viability Staining Solution (BioLegend, 420404). For intracellular DNA-PKcs flow, cells were stained using an intracellular flow cytometry kit (Cell Signaling, #13593S) according to the manufacturer’s instructions using anti-DNA-PKcs (Cell Signaling, #38168) and goat anti-rabbit IgG (Invitrogen, #A32733).

### Metabolic analysis

Extracellular acidification rate (ECAR) was measured using the Seahorse XFe bioanalyzer. Note that 2 × 10^5^ T cells per well (≥8 wells per sample) were spun onto Cell-Tak (Corning)–coated seahorse 96-well plates and preincubated at 37°C for approximately 20 minutes in the absence of CO_2_. ECAR was measured in XF media (nonbuffered RPMI 1640 containing 10 mmol/L glucose, 2 mmol/L l-glutamine, and 1 mmol/L sodium pyruvate) under basal conditions and in response to activation (SIINFEKL or anti-CD3/28), 2 μmol/L oligomycin, and 10 mmol/L 2-Deoxy-D-glucose.

### In vitro killing assays

Co-culture experiments were performed as previously reported.^39^ Briefly, T-cell killing assays were carried out using preactivated CD8^+^ T cells from OTI (tumor-specific) mice. Activated T cells were washed and labeled with Cell Trace violet to enable their subsequent discrimination from target cells. Target cells (MC38^SIINFEKL^) were plated for 16 hours prior to culturing with T cells. The cocultures were conducted at different target: effector (T:E) ratios for 16 hours. Target cell viability was then determined by the percentage of Annexin V and propidium iodide (PI) cells.

### Proliferation assays

Prior to activation OTI T cells were labeled with 1µM Cell Trace Violet dye (ThermoFisher) and then activated with 1µg/mL SIINFEKL peptide for 48hrs and previously described. FACS analysis and Flow Jo software was used to calculate the division index using proliferation modeling.

### Real-time PCR

Cells were lysed with Trizol (Qiagen, #15596026) and total RNA was isolated using Zymo Direct-zol RNA MiniPrep (Zymo, #R2052) following the manufacturer’s protocol. cDNA synthesis was performed using iScript cDNA Synthesis Kit (Bio-Rad, #1708841) according to manufacturer’s instructions. PowerUP SYBR Green master mix (AppliedBiosystems, #A25742) was used for real-time PCR detection with primers listed below. Reactions were performed in technical triplicate and ΔΔCt method was used to calculate relative expression of the target transcripts normalized to TBP transcript levels.

Mouse primers:

Tbp: FWR-5’-AGAACAATCCAGACTAGCAGCA, REV-5’-GGGAACTTCACATCACAGCTC

Gzmb: FWR-5’-TCACAAGGACCAGCTCTGTCCT, REV-5’-GTTGGGTTGTCACAGCATGG

Prf1: FWR-5’-GAGAAGACCTATCAGGACCA, REV-5’-AGCCTGTGGTAAGCATG

Eomes: FWR-5’-CCCCTATGGCTCAAATTCC, REV-5’-CCAGAACCACTTCCACGAA

IFN*γ*: FWR-5’-ACAGCAAGGCGAAAAAGGATG, REV-5’-TGGTGGACCACTCGGATGA

### Western Blot

Samples were lysed in 0.5% SDS RIPA buffer with protease inhibitors (Thermo Scientific, #78425) and phosphatase inhibitors (Roche, # 04906837001) followed by sonication. Samples were heated in LDS (lithium dodecyl sulfate) loading buffer then loaded into 4–12% bis-tris gels (Thermo Scientific #NW04122BOX) followed by transfer to a PVDF membrane. Membrane was probed for AKT, pAKT, DNA-PKcs and Gapdh. Imaging was performed with a GE ImageQuant LAS4000.

### Statistical analysis

Analysis of significance was done using standard t-test and expressed as the mean ± standard deviation. Assays were performed in triplicate. P ≤ 0.05 was considered significant.

## Author’s Contributions

Ana Azevedo-Pouly and Lauren Appell – Participated in research design, performance of research, data analysis, and writing of the paper, Lyle Burdine – Participated in research design, data analysis, and writing of the paper, Lora J. Rogers – performance of research and data analysis, Lauren C. Morehead – performance of research and data analysis, Melanie Barker - performance of research, animal management, Zachary Waldrip – performance of research, Brian Koss - participated in research design, performance of research, data analysis and writing of the paper, Marie Schluterman Burdine – participated in research design, performance of research, data analysis, writing of the paper, and obtaining funding.

